# Viral cross-linking and solid-phase purification enables discovery of ribonucleoprotein complexes on incoming RNA virus genomes

**DOI:** 10.1101/2020.04.08.032441

**Authors:** Byungil Kim, Sarah Arcos, Katherine Rothamel, Manuel Ascano

## Abstract

The initial interactions between incoming, pre-replicated RNA virus genomes and host protein factors are important in infection and immunity. Yet there are no current methods to study these crucial events. We established VIR-CLASP (VIRal Cross-Linking And Solid-phase Purification) to identify the primary viral RNA-host protein interactions. First, host cells are infected with 4SU-labeled RNA viruses and irradiated with 365 nm light to crosslink 4SU-labeled viral genomes and interacting proteins from host or virus. The cross-linked RBPs are purified by solid-phase reversible immobilization (SPRI) beads with protein denaturing buffers, and then identified by proteomics. With VIR-CLASP, only the incoming viral genomes are labeled with 4SU, so cross-linking events specifically occur between proteins and pre-replicated viral genomic RNA. Since solid-phase purification under protein-denaturing conditions is used to pull-down total RNA and cross-linked RBPs, this facilitates investigation of potentially all RNA viruses, regardless of RNA sequence. Preparation of 4SU-labeled virus takes ∼7 days and VIR-CLASP takes 1 day.

## Introduction

We developed a method to capture interactions between incoming viral RNA genomes and cellular proteins, as these events are crucial to both viral infection and host innate immunity. VIR-CLASP innovates on previous techniques such as RAP-MS^1^, TUX-MS^2^, and cRIC^3^ by specifically isolating interactions with pre-replicated viral genomes and by using sequence-independent methods to isolate RBP-RNA complexes. Since VIR-CLASP does not use sequence-dependent isolation of RBP-RNA complexes, this technique does not suffer from biases associated with changing probe hybridization efficiencies due to dynamic ribonucleoprotein assembly over time, experimental condition, or cellular context.

### Development of the protocol

VIR-CLASP identifies early viral-host interactions by using a 3-step strategy: first, viruses are propagated in the presence of 4SU; second, isolated 4SU-labeled viruses are infected into unlabeled host cells followed by ultraviolet (UV) irradiation with 365 nm light; and third, crosslinked viral RNA-protein complexes are purified by a modified solid-phase reversible immobilization method (CLASP, Figure 1). As we outline in detail below, VIR-CLASP is ideally suited for the identification of RNA binding proteins interacting with infecting viral genomic RNA.

**Fig. 1:**
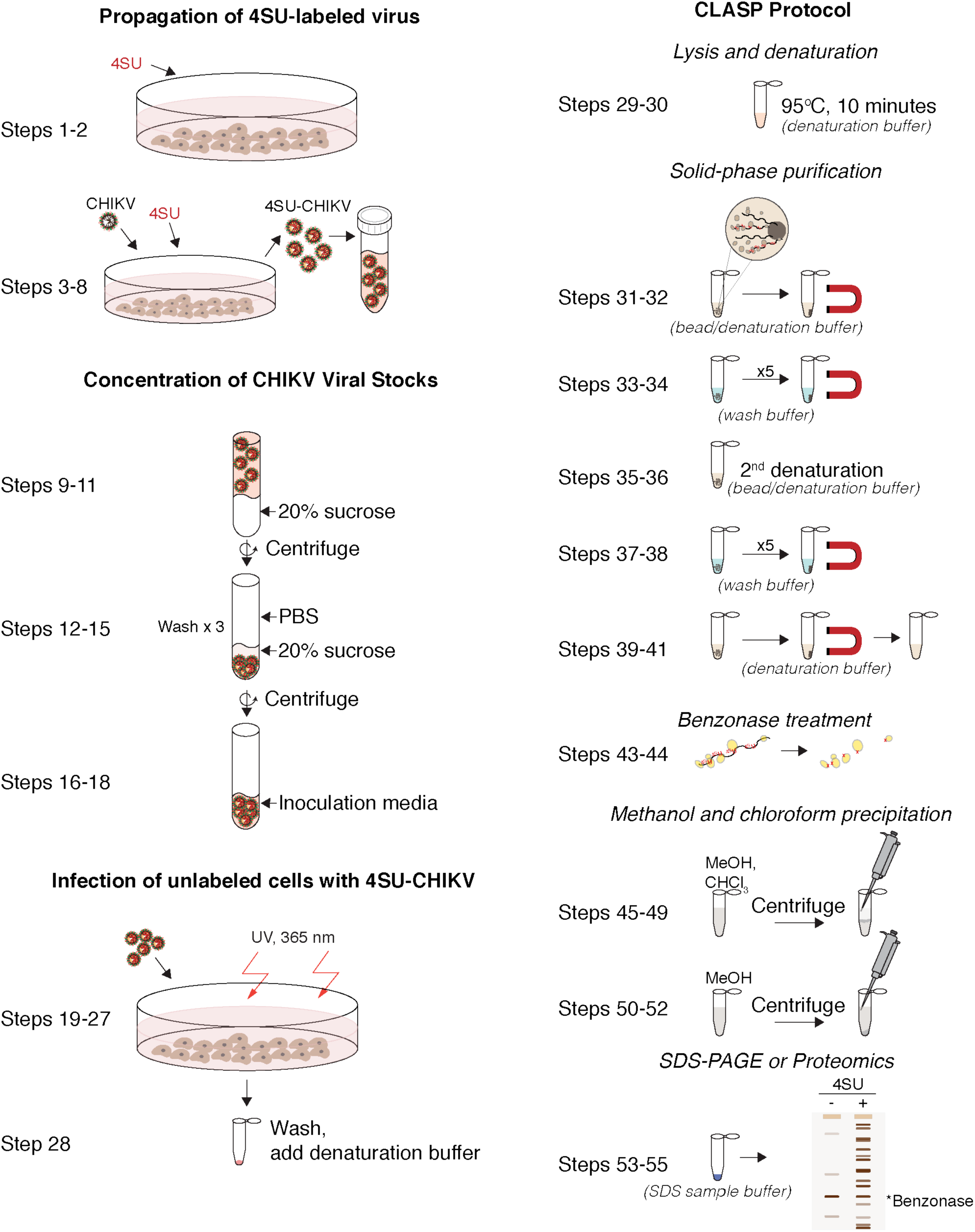
Outline of VIR-CLASP. VIR-CLASP is a method for identifying the primary interactions between viral RNA and host proteins. First, viruses are propagated with 4SU to label their RNA genome (Steps 1–18). Second, unlabeled cells are infected with 4SU labeled viruses (Steps 19–28). Finally, the RNP complexes are purified with SPRI beads under denaturing conditions and analyzed by Silver staining or LC–MS/MS (Steps 29-55). 4SU: 4-thiouridine, CHIKV: Chikungunya Virus, 4SU-CHIKV: 4SU labeled Chikungunya Virus, CLASP: Cross-Linking And Solid-phase Purification.

To label the viral genome with 4SU, RNA viruses are propagated during 4SU treatment and the virus is purified by ultracentrifugation with a sucrose cushion. For VIR-CLASP to specifically identify proteins bound to pre-replicated viral genomes, 4SU should only be present in the viral genome. To ensure this, we wash the pelleted virus to remove any remaining free 4SU in the viral supernatant. Then, cells are infected with un-labeled or 4SU-labeled viruses and any uninfected virus is washed away. The infected cells are irradiated with 365 nm UV light to activate 4SU, driving efficient RNA protein crosslinking^4^. Cells are lysed under denaturing buffer conditions and boiled in order to minimize non-specific interactions between proteins or between protein and RNA. To implement a method that could be applied to all RNA viruses in an unbiased way, we used solid-phase reversible immobilization (SPRI) beads for the purification of protein-RNA interactions. SPRI beads are used in established methods for purification and size-selection of DNA or RNA but have not yet been optimized for purification of protein cross-linked to RNA^5^. The purification under protein denaturing conditions is of total nucleic acids but only proteins that are covalently crosslinked to the RNA viral genomes containing 4SU are recovered. The SPRI beads and complexes are washed under the denaturing conditions used for the initial binding. The crosslinked RNA-protein complexes are eluted, the RNA is digested by nuclease (Benzonase), and then the proteins are identified by LC-MS/MS.

### Applications of the method

Our experience with VIR-CLASP indicates that it can be used for a wide variety of RNA viruses and is quick to implement because it uses a SPRI paramagnetic bead-based technology that does not require sequence-specific optimization to purify the viral RNA. This facilitates comparisons across time points, different cell lines, and different strains for the same virus. Owing to its reliance on photocrosslinks between RNA and protein, we anticipate that VIR-CLASP would be most effective for investigating the host protein interactome with the genomes of single-stranded RNA viruses. Double-stranded RNA viruses have the added complexity of intra-strand crosslinks, thus reducing the probability of RNA-protein photoadducts, as has also been observed for RNA-oriented crosslinking-immunoprecipitation (CLIP) techniques such as HITs-CLIP^6^, eCLIP^7^, iCLIP^8^, and PAR-CLIP^9,10^. We have performed initial VIR-CLASP experiments on viruses from seven distinct families, including plus- and minus-stranded RNA viruses: CHIKV (Chikungunya virus, *Togaviridae*), EMCV(Encephalomyocarditis virus, *Picornaviridae*), MHV (Mouse hepatitis virus, *Coronaviridae*), ZIKV (Zika virus, *Flaviviridae*), RVFV (Rift Valley Fever Virus, *Phenuiviridae*), IAV (Influenza A virus subtype H3N2, *Orthomyxoviridae*), and VSV (Vesicular stomatitis virus, *Rhabdoviridae*) (Kim and Arcos et al, in press). We have observed an array of distinct interactions between cellular proteins and 4SU-labeled viral genomes. We use VIR-CLASP to discover the earliest host protein-viral RNA interactions between human cells and Chikungunya virus (CHIKV) or Influenza A (IAV) virus. CHIKV has a positive-sense RNA genome and IAV has a negative sense RNA genome, thus these viruses likely form distinct interactions upon infection into host cells. Using VIR-CLASP followed by mass spectrometry, we identified hundreds of proteins that directly bind CHIKV and IAV. We found proteins that distinctly interact with CHIKV or IAV, and proteins that interact with both (Kim and Arcos et al, in press). VIR-CLASP will enhance our understanding of the early events in viral replication, opening new avenues for therapeutic innovation and prophylactic treatment of CHIKV and other RNA viruses. VIR-CLASP can also be used to broadly identify the RNA binding proteins interacting with different RNA species (e.g., viral RNA within virion or cellular RNA). To identify RBPs in the virion, VIR-CLASP can be performed on just purified virus, without infection into host cells. We observed that viral proteins E1, E2, and Capsid interact with the CHIKV genome (Kim and Arcos et al, in press). Finally, a subsection of the VIR-CLASP protocol is also amenable for the broad identification of cellular RNA binding proteins that directly interact with bulk cellular RNA; labeling of cultured cells with 4SU allows for the isolation of RNA binding proteins with the CLASP method, in a sequence-independent way (Kim and Arcos et al, in press). This application of CLASP adds to the growing list of interactome-wide RBP identification methods^11-15^.

### Comparison with other methods

VIR-CLASP differs from RNA-antisense purification mass-spectrometry (RAP-MS^1^), thiouracil cross-linking mass-spectrometry (TUX-MS^2^) and comparative RNA-interactome capture (cRIC^3^) in two fundamental ways: first, VIR-CLASP only identifies interactions with the pre-replicated viral genome; second, VIR-CLASP uses a sequence-independent method to purify total RNA under protein-denaturing conditions. With VIR-CLASP we can identify host proteins that interact with the pre-replicated genomes of various RNA viruses.

Recently, other non-sequence-based methods of crosslinking RBP complexes and purification have been developed. Acid guanidinium thiocyanate-phenol-chloroform extraction (TRIZOL)^15^ and Phenol Toluol extraction (PTex) methods are based on chemically driven liquid phase separation. Silica-based solid-phase extraction^13,16^ uses silica matrices to purify RBP-RNA complexes. Compared to these other methods, the CLASP method is a paramagnetic bead-based approach. We find that crosslinked RBPs are more easily and specifically separated from the samples when using magnetic beads.

### Experimental design

#### Control (Non-labeled virus)

VIR-CLASP should be performed with non-labeled virus and 4SU-labeled virus simultaneously. As a blank, we used the LC-MS/MS-identified proteins from the sample with non-labeled virus to define the candidate set of proteins.

#### 4SU incorporation into viral genomes

Because VIR-CLASP requires 4SU labeled RNA, the 4SU incorporation rate in viral genomes is important. Depending upon the cell lines and virus, 4SU concentration and treatment time should be considered. If the viral propagation cycle is fast, pre-treatment of cells with 4SU is helpful because it allows cells to take up and phosphorylate 4SU before viral RNA replication starts. If the propagation cell line has not been used for a 4SU-labeling experiment previously, we normally treat the cells with 100 µM 4SU overnight (16 h) and analyze the 4SU incorporation rate of cellular RNA to verify that this cell line can uptake 4SU. Although a higher 4SU concentration would result in more 4SU incorporation into viral genomes, it could also impact the yield of virus propagation. We observed that titers of viral supernatant were 2∼3 fold decreased with CHIKV and ∼10 fold dropped with IAV after treating with 1 mM 4SU in the propagation medium. We typically use 100 µM 4SU for 16 h to pretreat and 1 mM 4SU for 2 days during propagation. The average amount of 4SU incorporation in the purified CHIKV particles was measured to be 2.5% using HPLC (Kim and Arcos et al, in press).

#### Test the timing of pre-replication with titer assay and qPCR

To define the timepoints when interactions between pre-replicated virus genomes and host proteins are most important, a viral titer assay of UV-irradiated 4SU-labeled virus at timepoints before and during infection can be used. Covalent crosslinks between viral RNA and proteins inhibit maturation of new virions only when the incoming RNA genomes are necessary for critical steps including translation and replication. With 4SU-labeled CHIKV, we observed that the initial decrease in titer when irradiating infected cells at 0 hpi was from ∼10^9^ to 10^7^; therefore, we estimate that ∼99% of viral particles contain a biologically meaningful amount of 4SU substitution (Kim and Arcos et al, in press). The UV irradiated viral titers recover to about to control levels at 3 hpi. RT-qPCR analysis also revealed that CHIKV RNA increases between 2 and 3 hpi, confirming that viral transcription and replication have begun (Kim and Arcos et al, in press). Therefore, for CHIKV, we chose to examine pre-replicated viral RNA and host protein interactions between 0 hpi and 3 hpi. We observed that IAV replication is fully dependent upon the pre-replicated RNA genome at ∼ 1 hpi, and does not become independent until ∼ 4 hpi (Kim and Arcos et al, in press).

#### Determine the bead buffer ratio

The SPRI method uses carboxyl-coated magnetic beads that can reversibly bind RNA in SPRI bead buffer (20% polyethylene glycol (PEG) and 2.5M NaCl)^5^. By increasing the ratio of SPRI bead buffer volume to sample volume, the efficiency of binding smaller RNA will be increased^17^. In the VIR-CLASP method, we employed protein denaturing conditions and SPRI beads to purify crosslinked RNA-protein complexes. As in the SPRI method, the CLASP bead buffer ratio can affect the efficiency of binding smaller RNA. Although increasing the bead buffer ratio can purify smaller RNA, it may be not suitable for the CLASP method since the solubility of higher salt concentration with denaturing agents is limited. Based on the length of the RNA viral genome, the CLASP bead buffer ratio should be optimized. VIR-CLASP can use up to 1:1 ratio (x1 bead buffer ratio) of CLASP bead buffer to sample buffer, which can recover longer than 300 nt RNA (Supplementary Fig. 1). In VIR-CLASP with CHIKV, we used 2:3 ratio (x0.66 bead buffer ratio) to recover CHIKV viral RNA and crosslinked protein.

### Expertise needed to implement the protocol

A competent graduate student or postdoc can perform all procedures for VIR-CLASP. It is essential for investigators working with infectious viral pathogens to obtain the appropriate level of biological safety training required by their respective institutions, and to operate the protocol within approved and certified biosafety cabinets and facilities. Core facilities for proteomics and operation of the LC–MS/MS instrument are also required.

### Limitations

In the VIR-CLASP method, the yield of cross-linked protein is dependent on 4SU-labeled RNA that is released from infected virus. To yield enough signal, we typically infect with a viral MOI of 200 to 5000. If viral propagation is not efficient, total virus quantity might also be an issue. As with all proteome-wide screens, it is recommended that identified hits are validated. For viral nucleic acid-host protein interactions, it is recommended that validation experiments are performed at lower MOI (e.g. 0.01 to 1). It is possible that depending on the virus, small amounts of cellular RNA can be packaged within the virion, and thus contains 4SU. However, IAV and CHIKV contain either no or very little cellular RNA within the virion (IAV: < 3%^18^, CHIKV: < 1.3%^19^) (Kim and Arcos et al, in press). It may be necessary to assess the extent of cellular RNA that can carry-over using reverse-transcription followed by PCR or sequencing methods. It is also possible that host proteins from the propagation cell line are packaged within the viral particle. If this is a concern, we recommend performing mass-spec analysis on viral particles after purification to determine the extent of cellular protein carryover during propagation.

## Materials

### Biological materials

CAUTION: The cell lines used in your research should be regularly checked to ensure that they are authentic and that they are not infected with mycoplasma. Cell lines for viral propagation and infection should be chosen based on the RNA virus used.

Cell line for Chikungunya virus propagation: BHK-21 (ATCC, cat. no. CCL-10)

Cell line for host cells for Chikungunya viral infection: U2OS (ATCC, cat. no. HTB-96)

Virus: CHIKV strain 181/25 (provided by Terence S. Dermody, University of Pittsburgh School of Medicine)

CAUTION: Virus must be handled according to approved biosafety protocols and regulations set by each individual lab and institution. Improper handling of samples may lead to exposure to infectious pathogens.

## REAGENTS

DMEM (Gibco, cat. no. 11965-118)

L-Glutamine (Gibco, cat. no. A2916801)

HEPES (Gibco, cat. no. 15630080)

Sodium Pyruvate (Gibco, cat. no. 11360070)

MEM Non-Essential Amino Acids Solution (Gibco, cat. no. 11140050)

2-Mercaptoethanol (Gibco, cat. no. 21985023)

McCoy’s 5A (Modified) Medium (Gibco, cat. no. 16600082)

4-Thiouridine (Sigma, cat. no. T4509)

DMSO (sigma, cat. no. D2650)

PBS (National Diagnostics, cat. no. CL-253)

FBS (Peak Serum, cat. no. PS-FB2)

Tris-base (Sigma, cat. no. 93362)

HCl (Sigma, cat. no. H1758)

NaCl (Sigma, cat. no. S3014)

EDTA (Sigma, cat. no. E6758)

NaOH (Sigma, cat. no. S8045)

Sucrose (Sigma, cat. no. 84097)

SDS (Sigma, cat. no. 74255)

NP-40 (Sigma, cat. no. I8896)

Glycerol (Sigma, cat. no. G5516)

Sera-mag SpeedBeads (Fisher, cat. no. 09-981-123)

Tween 20 (Sigma, cat. no. P9416)

PEG-8000 (Sigma, cat. no. 89510)

Benzonase (Novagen, cat. no. 70746)

MgCl_2_ (Sigma, cat. no. M8266)

DTT (Sigma, cat. no. D9779)

Methanol (Sigma, cat. no. 34860)

Chloroform (Sigma, cat. no. C2432)

NuPAGE LDS Sample Buffer (4X) (Invitrogen, cat. no. NP0007)

NuPAGE Sample Reducing Agent (10X) (Invitrogen, cat. no. NP0004)

## EQUIPMENT

Floor Low Speed Centrifuge (Thermo Fisher, cat. no. Sorvall RC-3B Plus)

H-6000A 6 x 1000mL Swinging-Bucket Rotor (Thermo Fisher, cat. no. 11250)

H-6000A bucket adapter for 500-ml tubes (Thermo Fisher, cat. no. 00-444)

Ultracentrifuge (Beckman coulter, cat. no. Optima LE-80k)

SW 32 Ti Swinging-Bucket Rotor (Beckman coulter, cat. no. 369650)

Spectrolinker XL-1500 (Spectronics, cat. no. XL-1500A)

DynaMag-2 Magnet (Thermo Fisher, cat. no. 12321D)

Microcentrifuges (Eppendorf, cat. no. 5404000138)

Vortex-Genie 2 (Scientific Industries, cat. no. SI-0236)

Digital Dry Bath (Fisher Scientific, cat. no. 11-715-125D)

ThermoMixer (Eppendorf, cat. no. 5382000023)

### Plasticware

150 mm TC-treated Cell Culture Dish (Corning, cat. no. 353025)

500 mL PP Centrifuge Tubes (Corning, cat. no. 431123)

Cell Scraper (VWR, cat. no. 101093-452)

1.5ml-tubes (Denville, cat. no. C2170)

Thinwall Polypropylene Tube, 25×89mm (Beckman coulter, cat. no. 326823)

0.45 µm filter (Millipore, cat. no. SCHVU05RE)

## REAGENT SETUP

Virus diluent

Virus diluent is 10 mM HEPES and 1% (v/v) FBS in DMEM medium.

Complete DMEM

Complete DMEM is 10mM HEPES, 2 mM L-Glutamine, 1 mM Sodium Pyruvate, 1x

MEM Non-Essential Amino Acids, 50 µM 2-Mercaptoethanol and 10% (v/v) FBS in DMEM medium.

4SU stock solution

Prepare 4SU stock solution at a concentration of 1 M in DMSO.

CRITICAL: 4SU is light sensitive.

20% sucrose in TNE buffer

20% sucrose in TNE buffer is 50 mM Tris–HCl [pH 7.4], 100 mM NaCl, 0.1 mM EDTA and 20% (w/v) sucrose in water.

3x denaturation buffer

3x denaturation buffer is 150 mM Tris–HCl [pH 6.8], 30% (v/v) Glycerol, 7.5% (w/v) SDS and 2% NP-40 in water.

SPRI bead stock

Sera-mag SpeedBeads are provided as a concentration of 50 mg/ml. After wash with TE (10 mM Tris–HCl [pH 8.0] and 1 mM EDTA) buffer, SPRI beads are reconstituted as a concentration of 1 mg/ml in 10 mM Tris–HCl [pH 8.0], 1 mM EDTA, 18% (w/v) PEG-8000, 1 M NaCl and 0.055% (v/v) Tween-20 in water. Store at 4 °C for up to 1 month.

Beads buffer

Beads buffer is 2.5 M NaCl and 20% (w/v) PEG-8000 in water.

WASH buffer

WASH buffer is prepared as mixed with 1x denaturation buffer and Beads buffer in a 0.66x bead buffer ratio by volume.

4x Benzonase reaction buffer

4x Benzonase reaction buffer is 80 mM Tris–HCl [pH 7.5], 20 mM MgCl_2_, 600 mM NaCl, 40% (w/v) Glycerol and 4 mM DTT in water.

CRITICAL: DTT should be added freshly before use.

### Procedure

#### Propagation of 4SU-labeled virus

1. Seed 5×10^6^ BHK-21 cells in a 150 mm Cell Culture Dish in 20 ml of complete DMEM with or without 100 µM 4SU. We typically use 12 plates per condition.

2. Incubate cells at 37 °C overnight.

3. Remove medium from flask, and incubate cells with 2.5 ml of diluted inoculum of CHIKV as MOI 0.1, at 37 °C for 1 h with rocking every 10-15 min to prevent cells from drying.

4. After 1 h, remove inoculum and wash cells twice with 1X with PBS. CRITICAL STEP: Ensure complete washing of uninfected CHIKV that is not labeled with 4SU, as this could affect the measured 4SU incorporation rate.

5. Add 17 ml of complete DMEM with or without 1 mM 4SU to the cell culture dish, and incubate cells until cytopathic effects are observed (typically 48 h post-infection) at which point most of the cells will have lifted or rounded.

### TROUBLESHOOTING

6. After cells have rounded and lifted, harvest and transfer the viral supernatants to a 500 ml conical tube.

7. Clear cell debris by centrifugation at 1,000 x *g* for 10 min and filter through a 0.45 μm filter.

8. Transfer to a new conical, and aliquot.

CAUTION: Each aliquot is only to be thawed once to maintain an accurate titer. PAUSE POINT: The virus can be frozen at -80 °C until use.

### Concentration of 4SU labeled CHIKV

9. Transfer 30 ml of supernatant to each ultracentrifuge tube.

10. Underlay with 5 ml of sterile 20% sucrose in TNE buffer.

CRITICAL STEP: The concentration of the sucrose cushion should be adjusted based upon the density of the virion. The density of CHIKV is 1.22 g/cm^3^ in sucrose^20^.

11. Ultracentrifuge with an SW32Ti swinging bucket rotor at 121,000 × g for 3 h at 4 °C.

12. Remove supernatant. Pellet may or may not be visible, but virus will be stuck to bottom of tube.

13. To wash away free 4SU, add 30 ml of cold PBS on top of the pellet, and underlay with 5 ml of sterile 20% sucrose in TNE buffer.

14. Ultracentrifuge with an SW32Ti swinging bucket rotor at 121,000 × g for 30 min at 4 °C and remove supernatant.

15. Repeat Steps 13-14 two further times to completely rinse the free 4SU out of the virus pellet.

16. Add 1 ml (or less) of cold virus diluent to the pellet (do not pipette up and down yet).

17. Parafilm tubes and allow the virus to “soak” at 4 °C for 4-6 hours.

PAUSE POINT: The virus pellet can be “soaked” for up to 12 hours (overnight).

18. Mix the soaked viral pellet well by pipetting up and down many times (>20).

CRITICAL STEP: Ensure complete suspension of the viral pellet, otherwise titer estimation may be inaccurate.

### Infection of unlabeled cells with 4SU-CHIKV

19. Seed 3×10^6^ U2OS cells in a 150 mm cell culture dish in 20 ml of complete DMEM.

CRITICAL STEP: To ensure that the cells are ready at the same time as the concentrated virus, perform step 19 at the same time as virus concentration (steps 9-17). Therefore steps 18 and 22 will occur on the same day.

20. Incubate cells at 37 °C overnight.

21. Remove medium from dish and wash twice with 5ml cold PBS.

22. Remove PBS from dish and incubate cells with 2.5 ml of diluted inoculum of CHIKV at MOI 1000, a 4 °C for 1 h with rocking every 10-15 min to prevent cells from drying.

23. After 1 h, remove inoculum and wash twice with PBS.

24. Add 17 ml complete DMEM and incubate at 37 °C until the crosslinking time point.

25. Remove medium from dish and wash twice with 5ml cold PBS.

26. Remove PBS from dish and add 2.5 ml PBS.

27. Irradiate uncovered dish twice with 0.6 J/cm^2^ of 365 nm UV light in a Spectrolinker XL-1500 or similar device.

28. Scrape cells from the plate with a cell scraper, transfer and divide 600 µl each to four 1.5-ml tubes and add 300 µl 3x denaturation buffer per tube. Mix well by pipetting up and down.

PAUSE POINT: Unless you want to continue directly with cell lysis, collect the cells by centrifugation at 500 x g for 5 min at 4 °C and discard the supernatant. Freeze the cell pellet in liquid nitrogen and store at -80 °C. When ready to proceed with CLASP, you can then add 1x denaturation buffer directly into cell pellets (∼900 µl/10^6^ cells) and mix well by pipetting up and down.

### CLASP

#### Preparation of whole cell lysate and protein denaturation

29. Incubate at 95°C for 10 min. Save an aliquot (after boiling).

PAUSE POINT: Cell lysate can be stored at -20 °C and the protocol restarted on Step 29.

30. Incubate sample at RT for 10 min.

#### Solid-phase purification

31. Add 600 µl CLASP bead suspension to 900 µl denatured-lysates to result in a 0.66x bead buffer ratio by volume.

CRITICAL STEP: Before adding the CLASP bead suspension to samples, warm up the beads to RT.

TROUBLESHOOTING

32. Mix by pipetting 10 times and incubate samples at 25 °C for 10 min; then place tubes on magnetic stand. Allow the CLASP beads to pellet to the magnet. Settle times will vary, and may be as long as 3 minutes.

33. Remove and discard the supernatant. Add 1 ml WASH buffer.

34. Mix by vortexing (max. setting) and allow the CLASP beads to re-pellet by magnet. Repeat Steps 33 and 34 four times for a total of five washes.

TROUBLESHOOTING

35. Remove last supernatant. Add 600 µl 1x denaturation buffer, and mix by vortexing – then incubate at RT for 5 minutes.

36. Add 400 µl beads buffer and mix the total reaction volume by pipetting 10 times and incubate sample at 25 °C for 10 min, then place tubes on magnetic stand.

37. Remove and discard the supernatant. Add 1 ml WASH buffer.

38. Mix by vortexing (max. setting) and allow the CLASP beads to re-pellet by magnet. Repeat Steps 37 and 38 four times for a total of five washes.

39. Remove final WASH supernatant then add 200 µL 1x denaturation buffer. Mix the total reaction volume by vortexing and incubate at RT for 5 minutes.

40. Collect beads by placing on magnetic stand.

TROUBLESHOOTING

41. Transfer supernatant to new tube. Save an aliquot.

42. (optional) If you want to check the recovery of RNA by the CLASP method, treat with 1 mg/ml proteinase K for 2 h at 55 °C and purify the RNA with Trizol.

#### Benzonase treatment

43. Add Benzonase reaction buffer with Benzonase to the supernatant from step 41.

44. Incubate samples at 37 °C for 2 h.

#### Methanol and Chloroform Precipitation

(Method can be scaled up or down.)

45. Split 800 µl benzonase-treated samples into two tubes with equal volume (400 µl)

46. Add an equal volume of 100% MeOH to each sample (50% MeOH final). Vortex (max. setting) for 3 sec.

47. Add 100 μl Chloroform. Vortex (max. setting) for 3 sec.

48. Spin for 2 min at 15,000 x g.

49. Remove the water/MeOH mix on top of the interface, being careful not to disturb the interface. Often the precipitated proteins do not make a visibly white interface, and care should be taken not to disturb the interface.

TROUBLESHOOTING

50. Add 400 μl of MeOH. Vortex (max. setting) for 10 sec.

51. Spin for 3 min at 15,000 x g; the protein precipitate should now pellet to the bottom of the tube.

52. Pipette as much MeOH as possible from the tube without disturbing the pellet and briefly dry the pellets in a vacuum centrifuge for 5 min at RT.

TROUBLESHOOTING

53. Re-suspend the pellets in 20-50 µl of 1x SDS-PAGE sample buffer (diluted with H_2_O), fully supplemented with fresh DTT.

CRITICAL STEP: The amount of 1x SDS-PAGE sample buffer used to re-suspend pellets can vary across virus/cell types based on 4SU incorporation, crosslink efficiency, and protein precipitation. Based on our experience, 20-50 µl is a typical range.

54. Incubate 95°C for 5 min.

55. Proceed silver stain or proteomics analysis

TROUBLESHOOTING

56. To analyze the raw LC-MS/MS data, please refer to our Supplementary Methods section, *Proteomics Data Analysis*.

**Timing:**

Step 1-18, Propagation of 4SU-labeled virus: 4 d (∼4 h hands-on)

Step 19-28, Infection of unlabeled cells with 4SU-CHIKV: 1 d (∼2 h hands-on)

Step 29-30, Preparation of whole cell lysate and protein denaturation: 30 min

Step 31-42, Solid-phase purification: 3 h

Step 43-44, Benzonase treatment: 2 h

Step 45-54 Methanol and Chloroform Precipitation: 30 min

Step 55 Silver staining or peptide identification and quantification by LC–MS/MS: 3-5 d

Step 56 Preliminary data analysis: 1-2 d depending on number of samples

### Troubleshooting

Troubleshooting advice can be found in Table 1.

**Table 1.** Troubleshooting table.

| Step | Problem | Possible reason | Solution |
| --- | --- | --- | --- |
| 5 | virus titer of 4SU labeled virus was too low compared to unlabeled virus | 4SU concentration was too high | Required a test for optimal concentration of 4SU |
|  | 4SU incorporation rate in 4SU labeled virus was too low. | 4SU concentration was too low or the uptake of 4SU was not efficient in the propagation cell line | Required a test for optimal concentration of 4SU |
| 31 | beads aggregate | The concentration of nucleic acid is too high | Use fewer cells or increase the lysis volume and bead quantity |
| 33-34 | wash buffer precipitation | Temperature is lower than 25 °C or the wash buffer is not prepared fresh | Warm up the temperature of the buffer |
| 40 | beads remain clumped | Nucleic acid is not eluted as well | Incubate another 5 min at 37 °C |
| 52 | The size of precipitated pellets are not different between unlabeled and 4SU conditions. | Background proteins bound to the beads | Avoid beads aggregation and wash buffer precipitation |
|  | Low yield of precipitated protein in 4SU condition | Low 4SU incorporation rate in 4SU labeled virus | Required a test for optimal concentration of 4SU |
|  |  | Low amount of 4SU labeled RNA | Optimize infection conditions (MOI, time) |
|  |  | Inefficient cross-linking | Optimize cross-linking conditions |
| 53 | pellet is not solublized | overdried pellets | Take more care in drying pellets |

### Anticipated results

Analysis by SDS-PAGE and silver staining of VIR-CLASP samples is expected to show enrichment of proteins with unique banding patterns across conditions/timepoints of cells infected with 4SU-labeled CHIKV (Figure 2). In the -4SU samples, which represent cells infected with unlabeled CHIKV, crosslinked, and processed using VIR-CLASP, the purification is anticipated to yield little protein (Figure 2, lanes 5 and 9). Only one protein, Benzonase, is added to digest the RNAs after solid-phase purification, and this protein should show similar intensity across the samples on the silver stain (Figure 2). By VIR-CLASP, the amount of enriched and intact ribosomal RNA should be unaltered between conditions, and independent of whether a sample contained crosslinked complexes (Figure 2). VIR-CLASP samples can be validated by immunoblot with previously known interacting and non-interacting RBPs. ELAVL1 (HuR) is an RBP known to interact with the CHIKV 3’ UTR^21^ and the antiviral RBP IFIT1 binds only to RNA viruses with a 5’ cap-0 structure that CHIKV lacks (Figure 2)^22^.

**Fig. 2:**
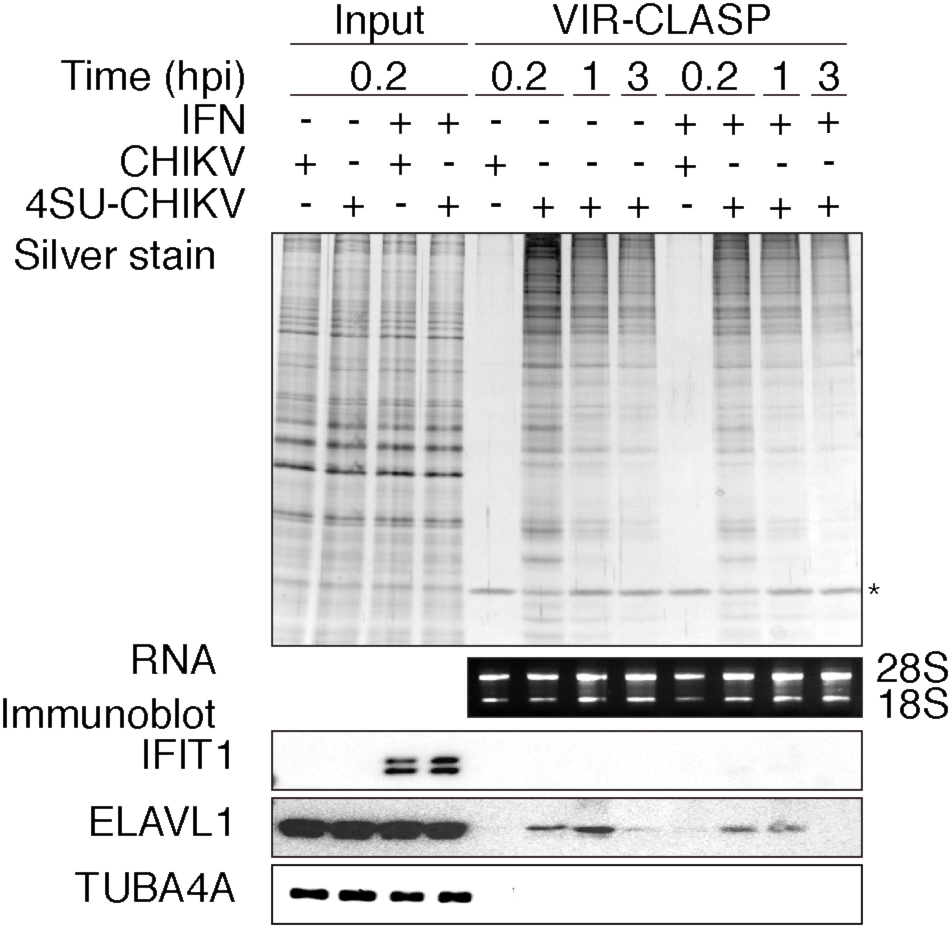
Anticipated results for VIR-CLASP samples. SDS-PAGE and silver stain (top), agarose gel (middle), or immunoblot (bottom) of proteins purified using VIR-CLASP from host (U2OS) cells that were infected with unlabeled or 4SU-labeled viruses (CHIKV). Condition (IFN): pre-treatment with recombinant interferon-β for 16 h before infection. Data represent three biologically independent repeats. Adapted with permission from [Kim and Arcos et al, in press], Cell Press.

The input to VIR-CLASP required in order to detect proteins with LC-MS/MS depends on the infectivity of the given virus in the given host cell. This is because only the pre-replicated viral genomes contain 4SU, so cellular protein content cannot be used as an input measure. For example, we observed that the strain of CHIKV used was much more efficient in deploying its genomic RNA into host cells than the strain of IAV used. This is reflected in the amount of viral and cellular input needed to get enough material to perform LC-MS/MS. For VIR-CLASP with CHIKV, infecting one 15 cm plate of U2OS cells (5 *10^6^) with MOI 1000 yielded enough purified protein to perform LC-MS/MS analysis at 0.2 hpi (Figure 2), but for IAV, we infected six 15 cm plates of A549 cells (6 * 10^7^ cells total) with MOI 1000 (Kim and Arcos et al, in press). For other viruses, optimization will be needed to yield sufficient protein for proteomics analysis.

We recommend performing a minimum of three biological replicates of VIR-CLASP for a given condition and virus, on different days. We also recommend that VIR-CLASP recovered eluates be subjected to LC-MS/MS analysis on the same day especially if label-free quantitation will be used.

To analyze the LC-MS/MS data, researchers can use their preferred software for peptide-spectrum matching. We advise including protein sequences for viral proteins as well as host proteins. For example, for CHIKV we identified the viral proteins Capsid, E1, and E2; we identified Nucleoprotein for IAV. To define candidate “VIR-CLASP RBPs” from the LC-MS/MS data peptide intensity ratios fo +4SU over -4SU samples can be calculated, and the average log2-transformed ratios for each protein can be tested against a null hypothesis of zero fold-change using the moderated t-test implemented in the R package limma^23^ (see Supplementary Methods) (Figure 3). A correction for multiple testing and a statistical significance cut-off should be applied. We found that applying the criteria of 0.01% FDR and fold change > 5 yielded ∼400 proteins significantly enriched in each VIR-CLASP condition for CHIKV. It should be noted that the biochemical stringency of the approach will likely yield very few recovered proteins in the -4SU sample. As a result, peptide intensity ratios cannot be calculated for all proteins because of the low complexity of the -4SU samples. For these proteins one can use a semiquantitative approach that assumes that peptides with zero intensity values are below the detection threshold^3,24^ (Figure 3). After determining the candidate “VIR-CLASP RBPs”, comparisons can be made between the conditions, timepoints, viruses, or cell lines tested. For example, overlaps between conditions for CHIKV showed that ∼255 candidate proteins interact with pre-replicated CHIKV RNA in all conditions tested and ∼340 proteins were unique to certain subsets of conditions (Figure 3). The choice of further functional analyses to perform will be dependent upon the particular experimental goals of the researchers.

**Fig. 3:**
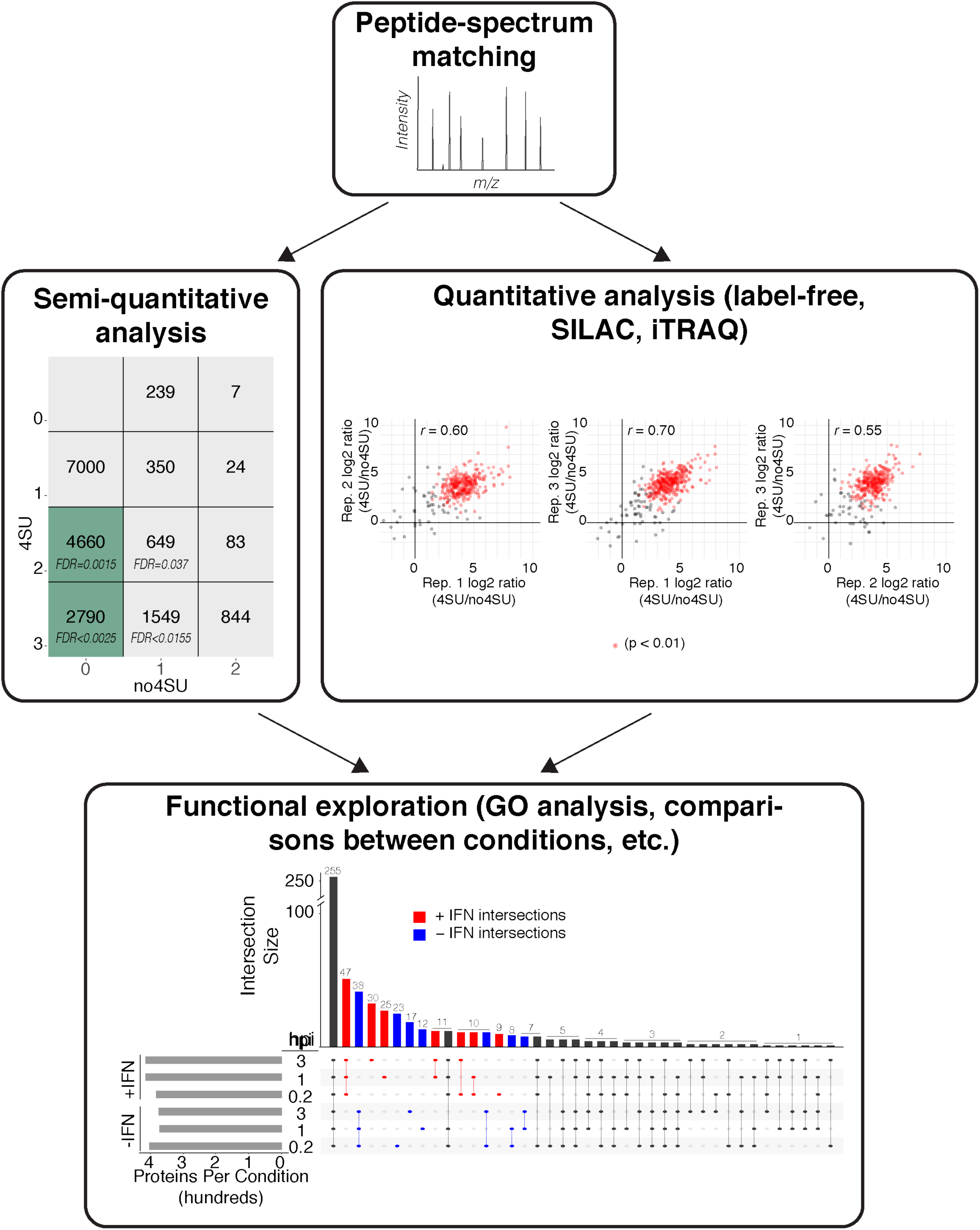
Proteomic Analysis of VIR-CLASP. Flow-chart illustrating an example computational analysis of VIR-CLASP performed with CHIKV. **Semiquantitative analysis** shows the matrix used to calculate FDRs for proteins which cannot be tested quantitatively due to the low complexity of the - 4SU samples. Peptides were sorted into boxes based on the number of biological replicates in which they were identified. Green boxes contain peptides with FDR < 0.01. **Quantitative analysis** shows the representative scatterplots of log2-intensity ratios of quantifiable proteins in +4SU over -4SU samples of VIR-CLASP with CHIKV (pairwise comparisons between three biological replicates). Red dots indicate significantly enriched proteins between all three biological replicates. **Functional analysis** shows an UpSet diagram^25^ summarizing the shared proteins that were identified in different timepoints and conditions of VIR-CLASP for CHIKV. The vertical bar chart shows the number of protein identifications common to the conditions marked by the colored and connected dots below. The horizontal bar chart shows the total count of proteins in each condition. Blue or red coloring highlights protein identifications unique to either (-) or (+) IFN pre-treatment. Adapted with permission from [Kim and Arcos et al, in press], Cell Press.

## Acknowledgements

We would like to thank Dr. Kristie L. Rose and Dr. W. Hayes McDonald at the Vanderbilt Mass Spectrometry Research Center for processing of the MS samples; Matthew Albertolle for technical help in the MS analysis; Dr. Thomas Voss and Dr. James E. Crowe Jr. (Vanderbilt University Medical Center) for IAV; and Dr. Terence S. Dermody (University of Pittsburgh School of Medicine) for CHIKV. Finally, we would like to thank members of the Ascano laboratory for their support, collegiality, and critical review of the manuscript. This work was supported by the National Institutes of Health 1R35GM119569-01 (M.A.), CTSA award No.UL1TR000445 from the National Center for Advancing Translational Sciences (B.K. and M.A.), Vanderbilt University Dept. Biochemistry start-up funds (M.A.), the Chemical Biology of Infectious Disease training grant 5T32AI11254-02 (S.A.), and the Chemistry-Biology Interface training grant 5T32GM065086-14.

## Author contributions

B.K. and M.A. designed, and B.K. optimized the VIR-CLASP technique. S.A. performed the proteomic analysis and data visualization. S.A., B.K., and M.A. wrote the paper.

## Supplementary Figure Legend

### Supplementary Figure 1: CLASP with RNA marker

Agarose gel of RNAs pull-downed from RNA marker using CLASP.

## Supplementary Methods

Proteomics Data Analysis

