## Supplementary Information for "Viral cross-linking and solid-phase purification enables discovery of ribonucleoprotein complexes on incoming RNA virus genomes"

### ARTICLE TITLE

### SUPPLEMENTARY METHOD – PROTEOMICS DATA ANALYSIS

#### ***Peptide-spectrum matching***

Raw data files from the LC-MS/MS instrument should be processed and searched with the researchers preferred peptide-spectrum matching software. In the case of VIR-CLASP with CHIKV and IAV, MaxQuant (v1.5.0.35)[1] was used to to generate peak lists and identify peptide-spectrum matches from Thermofisher raw files. The database of proteins sequences used should contain protein sequences from the host species as well as from the virus of interest. For CHIKV and IAV, searches were performed using a Uniprot/Swissprot database for *Homo sapiens* with only reviewed proteins included (downloaded on Feb. 28$^{th}$, 2018). Sequences were added to the database for Benzonase nuclease (Uniprot #P13717), for CHIKV proteins (Capsid, E1, E2, E3, 6k, nsP1, nsP2, nsP3, nsP4) from strain 181/25 (TSI-GSD-218), and for IAV proteins (H9XN78, 79, 80, 81, 83, 84, 85). For the purified CHIKV viral particles, we used a database containing the Uniprot reference proteome for *Mesocricetus auratus* (UP000189706- downloaded July 26$^{th}$, 2019, with one protein sequence per gene); sequences were added for CHIKV proteins (Capsid, E1, E2, E3, 6k, nsP1, nsP2, nsP3, nsP4) from strain 181/25 (TSI-GSD-218). Search parameters should be discussed and optimized with input from a proteomics or mass-spec expert. For CHIKV and IAV, the search parameters for Andromeda were: full tryptic specificity, two missed cleavages allowed, carbamidomethyl (C) fixed modification, and acetylation (N terminal) variable modification. Match between runs was selected. LFQ normalization was performed in separate parameter groups (+IFN and +4SU, -IFN and +4SU, +IFN and -4SU, and -IFN and -4SU). All other settings were left as default. This resulted in a protein FDR of < 1% for each dataset.

#### ***Quantitative analysis***

To define the set of candidate "VIR-CLASP RBPs", the peptide intensity ratios between +4SU and -4SU samples for proteins with a minimum of two distinct peptides can be calculated[2,3]. The average log2-intensity ratio for each protein can then be tested against a null hypothesis of 0 using a moderated t-test implemented in the R/bioconductor package limma[4]. p-values should be corrected for multiple testing (we used the Benjamini-Hochberg method). For CHIKV and IAV, proteins with a Benjamini-Hochberg adjusted p-value < 0.01 and a fold change greater than five were classified as candidate "VIR-CLASP RBPs". Correlations between replicates can be performed using Pearson correlation with the base R cor function, and the "pairwise.complete.obs" option to allow for missing values.

***Semi-quantitative analysis***

Due to the few proteins identified in the -4SU samples, peptide intensity ratios cannot be calculated for every protein. For proteins with zero intensity values in the -4SU samples, one can perform a semiquantitative approach that supposes that peptides with zero intensity values are below the detection threshold[2,3]. This method tabulates the number of replicates of +4SU and -4SU samples in which a peptide has a non-zero intensity value. For CHIKV, this produces a matrix of 12 different groups: peptides that were detected in 0, 1, 2, or 3 +4SU samples and 0, 1, or 2 -4SU samples (Figure 3B). For IAV, this matrix has 9 different groups: 2 replicates performed for +4SU and -4SU each. The FDRs were estimated as previously described[3], as the ratios resulting from dividing the transposed matrix. Using this semiquantitative approach, proteins were determined to be a "candidate VIR-CLASP RBP" for CHIKV or IAV if they comprised peptides found in cells that had an FDR < 0.01.

***Functional analysis***

Visualization of shared proteins between conditions can be achieved in R using the UpSetR package[5] for UpSet diagrams. Other functional analyses will be dependent upon the particular experimental questions asked.

All code for the R analysis will be made available at https://github.com/Ascano-Lab

### Supplementary Figure 1

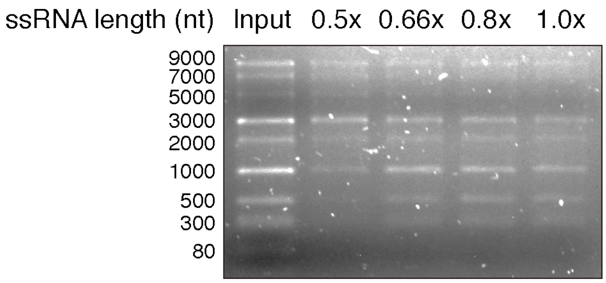
